# Msc1 is a nuclear envelope protein that reinforces DNA repair in late mitosis

**DOI:** 10.1101/2023.11.21.568106

**Authors:** Sara Medina-Suárez, Jessel Ayra-Plasencia, Lara Pérez-Martínez, Falk Butter, Félix Machín

## Abstract

Precise double-strand break (DSB) repair is paramount for genome stability. Homologous recombination (HR) repairs DSBs when cyclin-dependent kinase (CDK) activity is high, which correlates with the availability of the sister chromatid as a template. However, anaphase and telophase are paradoxical scenarios since high CDK favors HR despite sister chromatids being no longer aligned. To identify factors specifically involved in DSB repair in late mitosis, we have undertaken comparative proteomics in *Saccharomyces cerevisiae* and found that Msc1, a poorly characterized nuclear envelope protein, is significantly enriched upon both random and guided DSBs. We further show that Δ*msc1* is more sensitive to DSBs in late mitosis, and has a delayed repair of DBSs, as indicated by increased Rad53 hyperphosphorylation, fewer Rad52 repair factories, and slower HR completion. We discuss how Msc1 may favor the formation of Rad52 factories and the timely completion of HR before cytokinesis.

## Introduction

DNA double-strand breaks (DSBs) pose a threat for cell survival and genome stability, playing a major role in carcinogenesis ^1–3^. Cells deal with DSBs through two main DNA repair mechanisms, non-homologous end joining (NHEJ) and homologous recombination (HR). The former comprises error-prone pathways that join two broken DNA ends, whereas the latter involves pathways that use intact homologous sequences to restore the broken DNA sequence ^2,4–6^. The right choice of the DSB repair mechanism is crucial for the correct restoration of the DNA molecule. HR requires a well-aligned sister chromatid to be error-free, thus it is prioritized when DSBs occur in the S and G_2_ phases of the cell cycle. By contrast, NHEJ is favored in G_1_ phase, when HR would tend to use error-prone templates such as homologous chromosomes. Cells regulate the choice between NHEJ and HR primarily on the basis of cyclin-dependent kinase (CDK) activity, since this correlates well with the absence or presence of a sister chromatid. NHEJ is preferred in G_1_, exactly when CDK activity is low, whereas HR is ubiquitously upregulated by CDK as its activity rises from S phase ^7,8^. However, as cells transit through M phase, the relationship between CDK, HR and DSB repair becomes more obscure, despite CDK activity remaining relatively high until the telophase-G1 transition ^9–13^. Most complex eukaryotes, including animal and plant cells, undergo a mitotic cell division in which chromosomes condense to a large extent in early M phase (prophase-metaphase), concomitant with the resolution of the sister arms. The last chromosomal region to resolve is the centromere at the onset of anaphase, when chromosome segregation occurs. This massive condensation and early arm resolution appears to be incompatible with HR, which is largely inhibited ^6,9,11–15^. In contrast, in simple eukaryotes such as yeast and other fungi, chromosomes barely condense in early M phase (referred to here as G2/M) and are resolved as they segregate in anaphase by an unzipping centromere-to-telomere mechanism ^12^. As a result, sister chromatids remain aligned and suitable for HR until the anaphase onset ^16,17^.

DSB signaling and repair has been studied primarily in G1, S, G2 and early M (G2/M in yeast). Nevertheless, how cells respond to DSBs occurring in the window that spans from anaphase onset to the telophase-G1 transition is poorly known. In higher eukaryotes, this lack of knowledge stems from technical limitations to synchronize cells after the anaphase onset. However, this is not an inconvenience in the yeast *Saccharomyces cerevisiae*, in which cells can be stably arrested in late anaphase and telophase by means of conditional mutants for the mitotic exit network (MEN). The kinase Cdc15 is critical for MEN, and *cdc15* mutants arrest cells in a late anaphase/telophase stage with most sister chromatids apparently resolved and segregated ^18,19^. Hereafter, we will refer to this *cdc15* stage as late mitosis (late-M). We have previously used this arrest to question how cells respond to DSBs in a state of high CDK but segregated sister chromatids, and found that sister chromatids partly reverse their segregation, allowing for *de novo* sister chromatid alignments that may serve for error-free HR repair ^20^. Accordingly, mutants for HR were as hypersensitive to DSBs in late-M as in G2/M.

In the present work, we sought to identify specific determinants of the DSB response in late-M that may differ from those previously reported in G2/M. To this end, we used comparative abundance proteomics and identified a small set of proteins whose levels are increased upon DSBs in late-M relative to G2/M. Among these, we found the poorly characterized meiotic sister chromatid 1 (Msc1) protein, originally reported to be important for channeling meiotic HR towards the homologous chromosome rather than the sister chromatid ^21^. We confirmed proteomics results by both Western blotting and microscopy, and genetically demonstrate that Msc1 particularly enhances the DSB repair in late-M. Importantly, we show that Msc1 negatively regulates hypersignaling of DSBs and positively regulates the formation of Rad52 factories, which establishes a novel regulatory player of DSB repair in late-M.

## Results

### Abundance proteomics identifies Msc1 as a protein significantly enriched after DSBs in late mitosis

To screen for proteins that may be specifically involved in DSB signaling and repair at late stages of the mitotic cycle, we designed an experimental setup whereby we compared the proteomes of cells blocked in G2/M and late-M and subjected to DSBs (Fig S1). To further strengthen the DSB specificity of proteome changes, we used two distinct sources of DSBs. On the one hand, we treated cells with phleomycin, a radiomimetic drug that generates multiple randomly located DSBs ^22^. Phleomycin intercalates into DNA and locally generates reactive oxygen species (ROS) that attack and chemically modify the DNA until it breaks apart. On the other hand, we have introduced a genetic modification in the tested strain that allows the inducible expression of the HO endonuclease, which generates one DSB at the *MAT* locus. In late-M, the number of DSBs is two because both sister chromatids are expected to be cut by HO. The expression of HO was driven by the newly developed β-estradiol promoter ^23^, a tight promoter that bypasses the need to change the growth media for expression. In each cell cycle stage, we paired the DSB treatment with the corresponding mock treatment (Fig 1A). In this way, we filtered out changes in protein levels due to DSB-independent factors, such as those needed for the cell cycle arrests.

**Figure 1.**
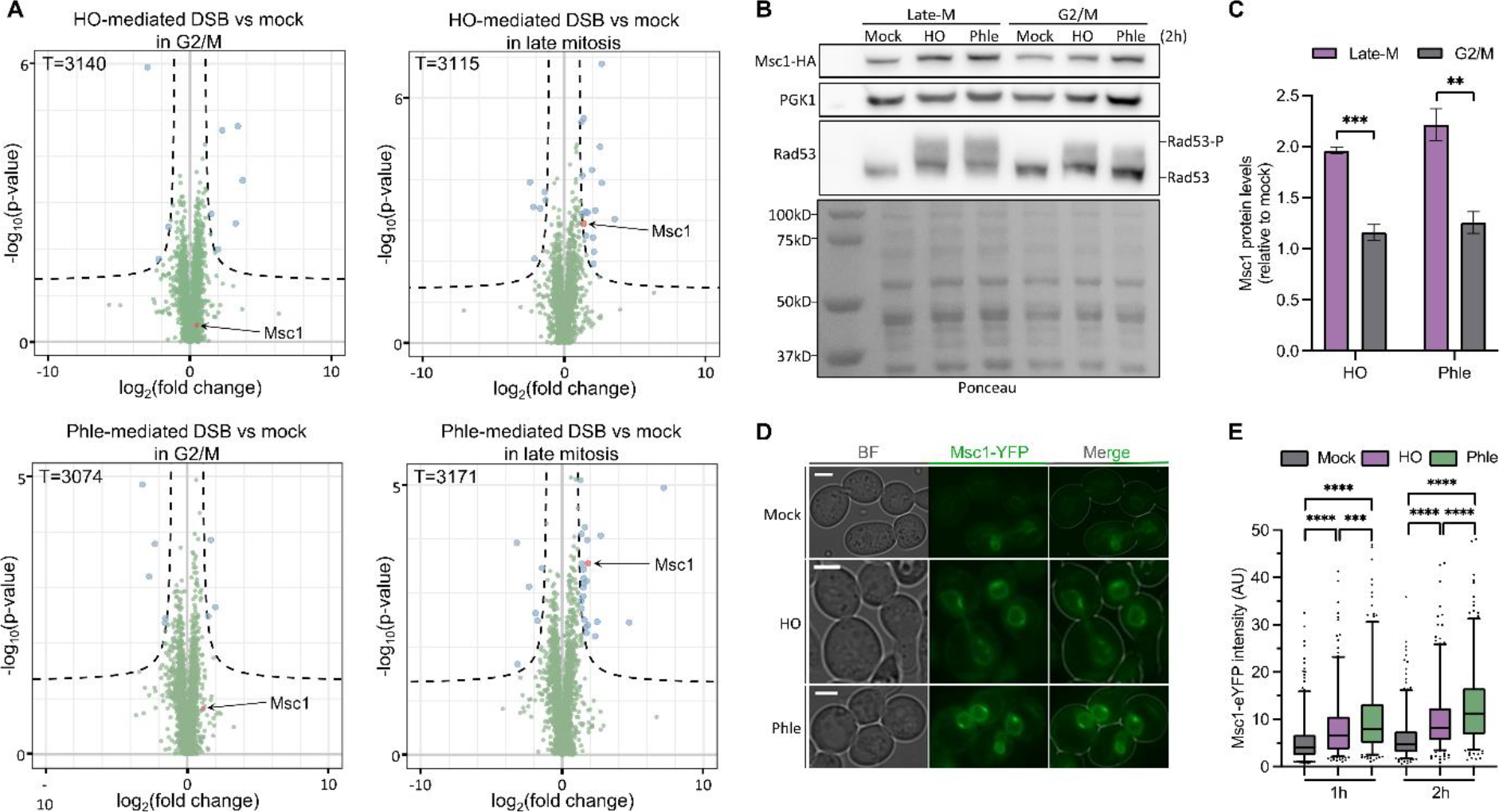
Proteomics for DSBs and Msc1 levels in G2/M and late mitosis. The experimental setup is schematized in Figure S1. **(A)** Volcano plots showing the fold change after DNA damage. For each cell cycle phase, the proteomic results of each type of DSB generated (HO and phleomycin) are compared separately with the mock control. Blue dots, enriched or depleted proteins; green dots, background proteins; red dot, Msc1. Total (T) detected proteins is indicated for each plot as well. **(B)**Western blot to confirm the Msc1 proteomics results. Ponceau staining of the membrane after transfer is also included. **(C)**Quantification of Msc1 after generating DSBs (relative to mock; mean ± s.e.m., n=3). The statistical analysis was performed by unpaired t test. **(D)**Representative micrographs of Msc1-eYFP in the mock experiment, HO- and phleomycin-mediated DSBs; 2 h after DSB induction. BF, bright field. **(E)**Quantification of Msc1-eYFP nuclear envelope levels in late mitosis, 1 and 2 h after DSB generation (mock, HO, and phleomycin). The box plots correspond to the pull of three independent experiments; >100 cells were quantified in each condition and experiment. Mann-Whitney tests were used for statistical significance.

The comparative proteomes revealed a number of proteins that were significantly enriched after DSB generation in both G2/M and late-M (Fig 1A; Table S1; Suppl excel file 1). Late-M resulted in more significant changes than G2/M, and the effect of phleomycin was stronger than that of HO induction. It is important to note that phleomycin is expected to modify the proteome in three ways: i) as a DSB generator itself, ii) as a ROS producer, and iii) as a xenobiotic. The products of five genes were specifically enriched in late-M after DSB generation with both phleomycin and HO; namely, *TFS1* (*YLR178C*), *GPH1* (*YPR160W*), *GND2* (*YGR256W*), *GDB1* (*YPR184W*) and *MSC1* (*YML128C*). Of these five, Msc1 is the only one that has been previously implicated in DSB signaling and repair, specifically in the choice between sister chromatids and homologous chromosomes during meiotic HR ^21^. None of the well-established factors involved in DSB signaling and repair were significantly enriched in any condition (Table S2; Suppl excel file 1). This suggests that constitutive levels of DSB proteins are sufficient to cope with DSBs, and this interpretation correlates well with previous transcriptomic data in which mRNA levels of these proteins changed little after DNA damage ^24^. Alternatively, any increase could be masked by post-translational modifications of DSB proteins that would affect their detection by mass spectrometry (e.g., phosphorylation, ubiquitination, sumoylation, etc.). Accordingly, our proteomics analysis detected about half of the proteins encoded in the yeast genome (∼3,000 proteins), but DSB repair proteins appeared clearly underrepresented (Table S2; Suppl excel file 1). Since Msc1 had been linked to HR before, we decided to focus on this particular protein for the rest of this work.

### Msc1 is nuclear envelope protein whose levels increase in late mitosis after DSBs

We began by validating the proteomic data that pointed towards a 4-fold increase of Msc1 in late-M after DSBs (2-fold increase after log2 transformation). To do this, we tagged Msc1 at the C-terminus with the HA epitope and measured Msc1 levels by Western blotting (Fig 1B,C). In these Western blots, we also included a housekeeping control for normalization, Pgk1, as well as the DSB sensor Rad53 as a reporter since it becomes hyperphosphorylated after DSB generation ^25^. Msc1 was enriched twofold after either phleomycin or HO treatments when compared to mock treatments. This enrichment was late-M specific as Msc1 barely changed after DSBs in the G2/M arrest.

Next, we investigated whether the increase of Msc1 levels was post-translationally regulated or as a result of an increase in *MSC1* expression. Hence, we measured *MSC1* mRNA levels by RT-qPCR in all tested conditions (Fig S2). We found that mRNA levels increased slightly after DSB generation, although neither the increase was that significant nor the cell cycle specificity that remarkable.

In addition to Western blots, we decided to address Msc1 levels by microscopy, which can report on the protein location and any shift that may occur after DSB generation (Fig 1D,E; S3). We tagged Msc1 with eYFP and found it at the nuclear periphery (nuclear envelope and/or perinuclear endoplasmic reticulum) in asynchronous and synchronized populations (Fig 1D and S3A). Interestingly, Msc1-eYFP abundance at the single cell level appeared to be highly variable, spanning up to 30-fold in NE intensity (Fig 1E, mocks), with cells where Msc1 was barely visible and cells with a very intense NE signal (Fig 1D and S3B). Both phleomycin and HO treatments led to a steady increase in Msc1-eYFP abundance (Fig 1D,E), until reaching, for example, a 4-fold change after 2h in phleomycin.

### Cells lacking Msc1 are hypersensitive to DSBs

Having validated that Msc1 levels increase after DSBs in late mitosis, we next addressed whether cells deficient in this protein are more sensitive to DSBs. Sensitivity tests based on spot assay experiments showed that the Δ*msc1* knockout mutant was more sensitive to phleomycin than its isogenic wild-type (WT) counterpart (Fig 2A). Similarly, Δ*msc1* was also hypersensitive to the DSB generated by the HO endonuclease (Fig 2B). For phleomycin, a similar hypersensitive profile was obtained in the growth curves (Fig 2C). Lastly, the Δ*msc1* strain was also hypersensitive to other forms of DNA insults that do not initially generate DSBs, although they do in the long term such as the replication stress drugs hydroxyurea (HU) and methyl methanesulfonate (MMS) (Fig S4A). In contrast, the mutant did not confer hypersensitivity to oxidative stress (Fig S4B). The latter profile reinforces that the phleomycin hypersensitivity is directly due to DSB formation and not to the concomitant production of ROS.

**Figure 2.**
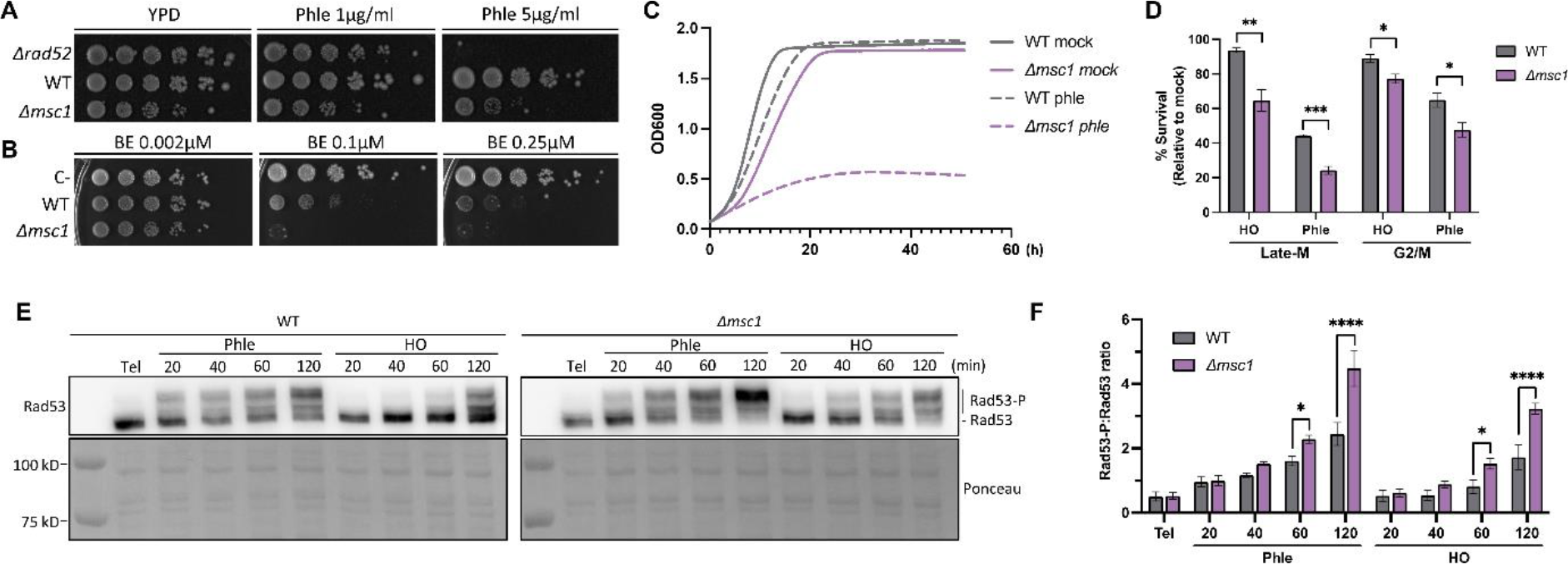
Sensitivity of *Δmsc1* to DNA damage. **(A)**Spot assay on phleomycin. A *rad52*Δ strain served as a positive control for sensitivity to DSBs. **(B)**Spot assay for HO DSBs. Both WT and *Δmsc1* strains carry the HO endonuclease under the control of the β-estradiol (BE) promoter. The *rad52*Δ strain used in (A) served here as a negative control (C-) as it does not carry the BE-HO system. **(C)**Growth curves of WT and *Δmsc1* with and without phleomycin (n=2). The s.e.m. is not shown for clarity (its highest value was less than 0.1). **(D)**Late-M versus G2/M clonogenic survival of WT and *Δmsc1* after DSBs (mean ± s.e.m., n=3). Survival was estimated relative to mock treatments (unpaired t-test). **(E)**Western blots of Rad53 hyperphosphorylation after DSBs in the WT and the *Δmsc1* strains. Tel, telophase (i.e., late mitosis). **(F)**Quantification of the Rad53-P:Rad53 ratio over the time course (mean ± s.e.m., n=3; unpaired t-test).

Although spot assays and growth curves clearly demonstrated the hypersensitivity of Δ*msc1* to DSBs, they could not discriminate whether this sensitivity was cell cycle specific, particularly whether or not late-M Δ*msc1* cells were more sensitive than their G2/M counterparts. To address this, we proceeded as shown in Fig S1 and determined the percentage of cells surviving DSBs in late-M and G2/M through clonogenic assays (Fig 2D). Relative to mock, the decrease in viability in the WT after HO induction was only 10% in both G2/M and late-M, whereas it was 40% and 60% in phleomycin, respectively. In the case of Δ*msc1*, these decreases were higher, and more severe in late-M than in G2/M (2-fold less viability in Δ*msc1* than in the WT in late-M, versus just 1.3-fold in G2/M).

### DNA damage signaling is higher in Δ*msc1*

To elucidate the molecular basis of the DSB hypersensitivity in Δ*msc1*, we first determined the kinetics of the DNA damage checkpoint (DDC). Rad53 is a master kinase in the DDC that is found hypophosphorylated in cells that are not experiencing DNA damage ^25^. By contrast, in the presence of ongoing DNA damage, including DSBs, Rad53 becomes hyperphosphorylated, and this molecular change is easily detected by Western blotting as an electrophoretic shift and the appearance of multiple slow-migrating Rad53 bands. Cells blocked in late-M prior to DSB generation showed a hypophosphorylated band (Fig 2E) ^20^, and once cells were either exposed to either phleomycin or HO expression, Rad53 became hyperphosphorylated. When we compared the kinetics of Rad53 phosphorylation in the WT and the Δ*msc1*, we observed that Rad53 became more hyperphosphorylated in the latter (Fig 2E,F), especially at the later time points of the experiment (1h and 2h).

To check whether this increase was due to a deficiency in DDC shutdown in Δ*msc1*, we repeated the DSB generation by HO induction, but washed out β-estradiol after 1h, thus allowing late-M cells to recover from the DSB insult. HO is known to be rather unstable and is rapidly degraded once the HO promoter is silenced ^26–28^. Rad53 remained hyperphosphorylated for the first 3 h after recovery and was gradually dephosphorylated over the next 6 h (Fig S5). By 18 h after recovery, Rad53 had reached the hypophosphorylated state seen before DSBs (or in the parallel mock treatments). Relative to the WT, no apparent delay in dephosphorylation kinetics was observed in Δ*msc1*.

### DSB repair by homologous recombination is slower in Δ*msc1*

The next issue we addressed was the kinetics of the DSB repair. To do this, we made use of the MAT switching system, a well-established reporter that allows both quantification of the DSB repair process and how much of it goes through either HR or NHEJ (Fig 3A and supplemental text) ^29^. In G2/M, the HO DSB is efficiently repaired through HR ^16,17^. Importantly, the HO DSB is also repaired by HR in late mitosis ^28^. To determine whether Msc1 impinges on the overall repair of the HO DSB, we compared the WT and the Δ*msc1* strains (Fig 3B,C). In both strains, most cells have efficiently cut the *MAT*a sequence after just 1 h of HO expression. Upon removal of the HO, the DSB began to be repaired by HR, yielding the *MAT*α product in more than 80% of the cases by 2 h. There was no difference between the WT and the Δ*msc1* mutant at this time; however, during the first 1.5 h, there was a clear delay in the Δ*msc1* (Fig 3C). This points out that Msc1 ensures an early repair of DSBs by HR, which may be critical for fast-cycling cells such as these, especially during the rapid transition through mitosis. No signs of repair through NHEJ were observed; however, to finetune whether any DSB could be channeled towards NHEJ, we measured the *MAT*a band throughout the experiment in derivative strains in which the *RAD52* gene had been deleted (Fig S6). Rad52 is an essential HR player ^30^, and gene conversion to *MAT*α is fully dependent on Rad52 ^31^. NHEJ was absent in late-M in both *MSC1* Δ*rad52* and Δ*msc1* Δ*rad52* (Fig S6).

**Figure 3.**
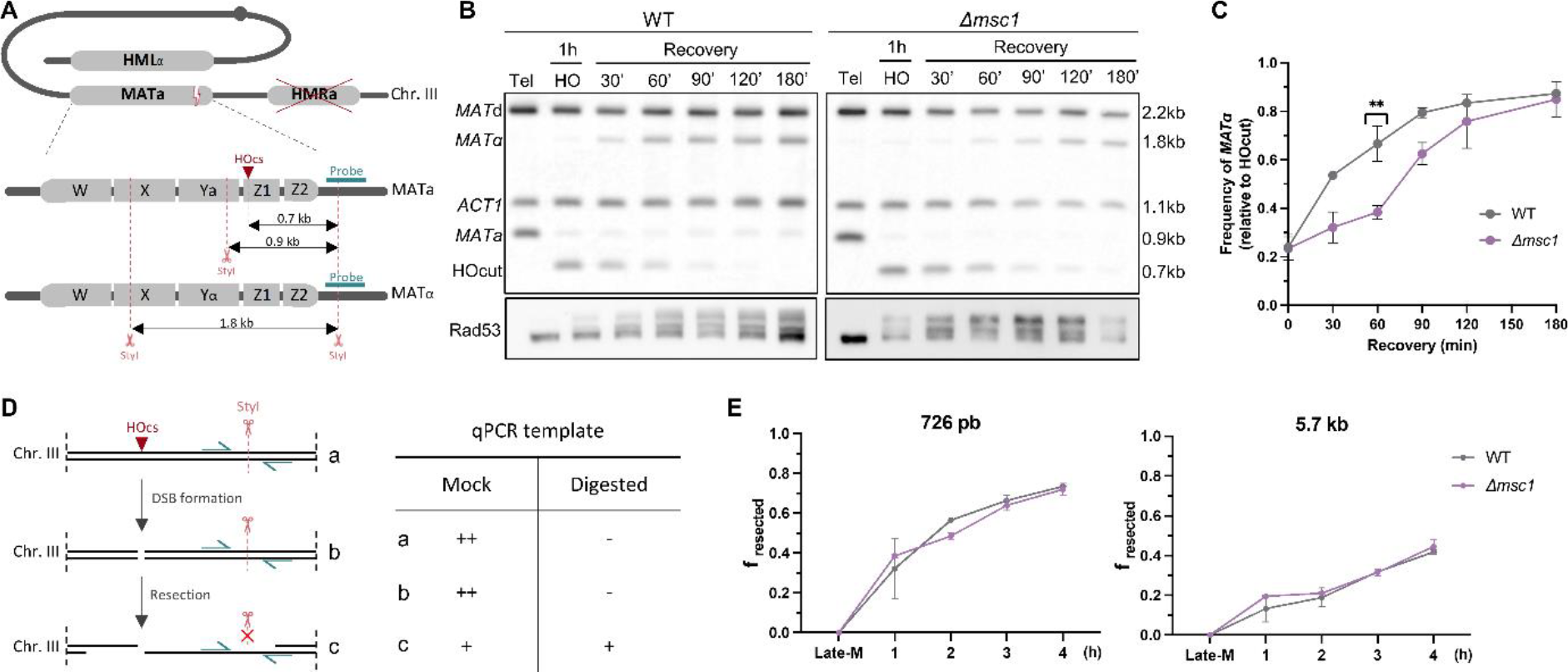
Late mitotic repair of the HO-mediated DSB in the WT and the *Δmsc1* strains. **(A)** Schematic of the *MAT* switch system used in this assay. On the top, the bent arrangement of chromosome III for intramolecular HR between the *MAT*a and the HMLα loci is shown. The position of the DSB in the *MAT*a locus is indicated by a dash; red crossed crosses indicate that the alternative recombinogenic *HMR*a locus is deleted. At the bottom, a zoomed view of the *MAT* alleles is depicted with the approximate location of the StyI restriction sites. Cutting with StyI differentiates the MATa and MATα alleles by Southern blot (fragment sizes are indicated by lines with double arrowheads; the probe is shown in blue). **(B)**Representative Southern blots of the MAT switch assay in the WT and *Δmsc1* strains in late-M. The Rad53 Western blot of the experiment is shown below. Tel, telophase (i.e., late mitosis). **(C)**Quantification of the *MAT*a conversion into *MAT*α (mean ± s.e.m., n=3). The switch was normalized to the amount of the *MAT*a cut after HO induction (the HOcut band). The unpaired t-test at 60’ is shown. **(D)**Principle of the qPCR assay used to measure resection at the HO cutting site (HOcs). On the left, schematic of the HOcs resection and its effects on PCR amplification. Primers (blue arrows) are designed to amplify sequences containing targets for StyI cleavage. On the right, a summary table of the expected amplification yield (a) before the HO cut, (b) after the HO cut but with resection not reaching the StyI site, and (c) with resection extending beyond the StyI site. Mock, no StyI digestion; Digested, StyI digestion; ++, extensive amplification, +, moderate amplification, - no amplification. **(E)**Resection kinetics for amplicons located at 726 bp and 5.7 bp downstream of the HO-generated DSB (mean ± s.e.m., n=2); f_resected_ is the fraction of resected DNA.

In order to channel DSBs for HR repair, DSB ends must first be resected, this is, processed into single-stranded DNA (ssDNA) so that these ssDNA tracts can search for homology in other genomic regions ^32^. To test whether resection was affected in Δ*msc1*, we compared resection efficiency at positions proximal (726 bp) and distal (5.7 Kb) to the HO DSB (Fig 3D,E). We measured resection based on a qPCR strategy capable of detecting ssDNA as it becomes resistant to *StyI* digestion (Fig 3D). For this experiment, we maintained HO expression throughout, observing that proximal and distal resection frequencies were equivalent in both the WT and the Δ*msc1* mutant (Fig 3E); as expected, proximal resection was faster. In *rad52*Δ, resection appears to be even more efficient (Fig S7), probably because of the inhibitory role of Rad52 on part of the resection machinery ^33^. Importantly, however, there was no difference between *MSC1* Δ*rad52* and Δ*msc1* Δ*rad52*.

### Msc1-deficient cells contain fewer Rad52 repair factories after DSBs

Having shown that DSB resection is not altered in Δ*msc1*, yet HR products are only obtained at later time points, we turned our attention to HR events that occur downstream. Resected DSBs are eventually coated by HR proteins involved in the search for homologous sequences and the formation of HR intermediates ^5^. All of these processes occur at distinct sites in the nucleus, which are referred to as DNA repair factories and where these HR factors are concentrated. These factories can be visualized by tagging HR proteins with fluorescent proteins ^34^. The most widely used representative of these factories is Rad52; thus, we followed Rad52-mCherry in late-M before and after DSBs (Fig 4A,B). We found that around 10% of cells arrested in the *cdc15-2* late-M presented Rad52 foci prior to DSB generation, and these values did not change during the mock treatments (Fig 4A). The percentages were equal for the WT and the Δ*msc1* strain. In the WT, HO-generated DSBs increased this percentage to ∼25%, whereas in phleomycin this percentage was even higher (∼45%). Interestingly, the amount of Δ*msc1* cells with Rad52 foci was significantly lower in both DSB scenarios, especially 1 h after DSB generation (∼15% and ∼20%, respectively). After 2 h, the fractions of Δ*msc1* cells with foci approached the WT values.

**Figure 4.**
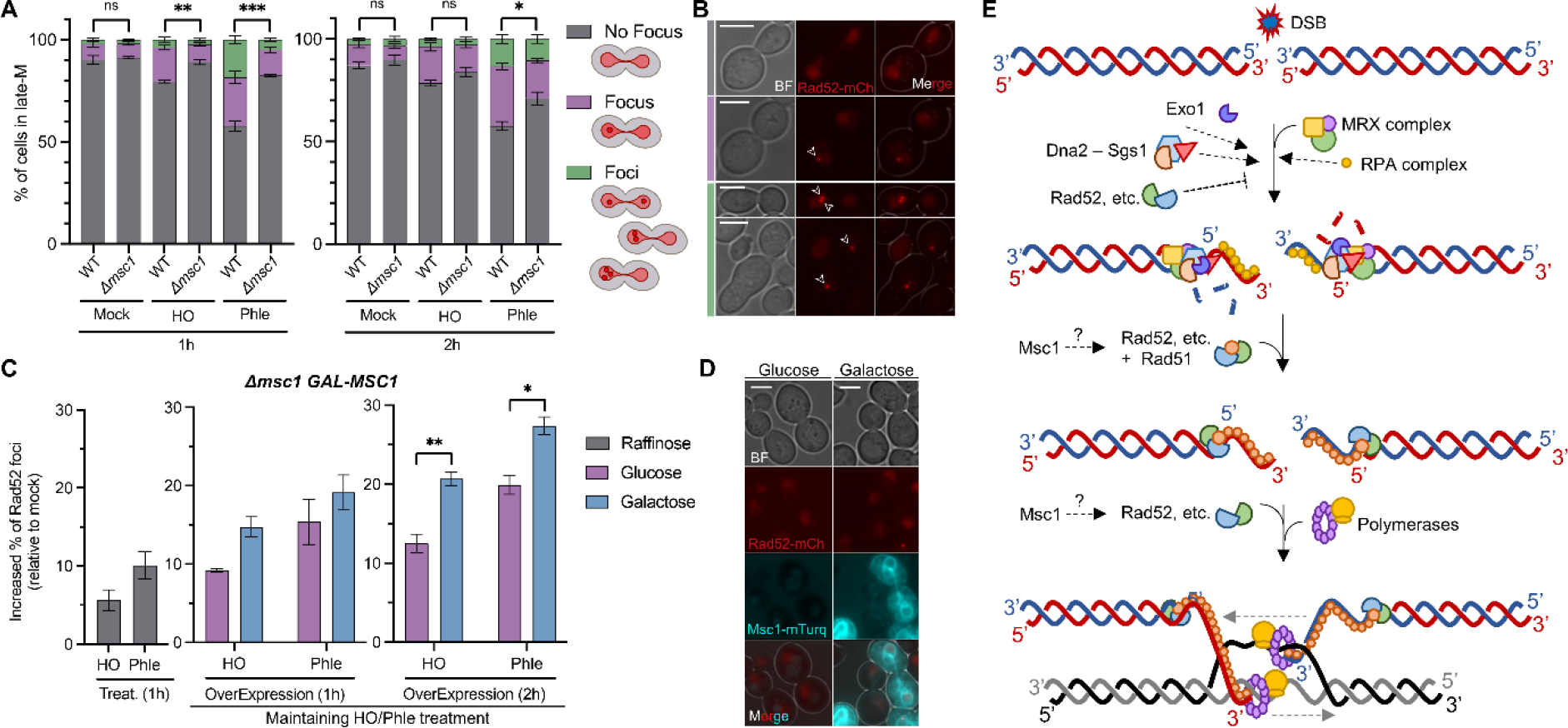
Msc1 affects the number of Rad52 repair factories in late mitosis. **(A)**Quantification of the absence of presence of late-M Rad52 foci after DSBs (mean ± s.e.m., n=3; unpaired t test). **(B)**Representative micrographs of late-M cells for each of the categories shown in (A). Micrographs in the same color pattern (shown as a vertical line on the left). White arrows point to Rad52 foci. **(C)**Effect of Msc1 overexpression on Rad52 foci after DSBs (mean ± s.e.m., n=3; unpaired t test). On the left (grey bars), the increase of late-M cells with Rad52 foci 1 h after DSB generation (relative to mock treatments); in the middle, the Rad52 foci increase 1 h after *MSC1* overexpression (and 2 h from DSBs); on the right, the Rad52 foci increase 2 h after *MSC1* overexpression (and 3 h from DSBs). The control subcultures without overexpression (glucose) are also shown. **(D)**Representative micrographs of late-M cells with (galactose) or without (glucose) Msc1-mTurquoise2. The examples correspond to the mock subcultures, 2 h after carbon source shift. BF, bright field. **(E)**Schematic of the putative position of Msc1 in the HR pathway. The 5’ ends of DSBs are recognized and resected by MRX, Exo1 and Dna2-Sgs1 and then coated by RPA. Rad52 modulates the resection, the replacement of RPA by Rad51 and the invasion of the template donor. Msc1 enhances a step between resection and template search.

Seeking to strengthen a functional relationship between Msc1 and Rad52, we next tested whether Msc1 overexpression could increase the number of Rad52 foci after DSBs in the Δ*msc1* background. For this purpose, we constructed strains where *MSC1* was under the control of the *GAL1-10* promoter (Fig S8A). This is a strong and inducible promoter which can be rapidly switched on by shifting the carbon source of the growth medium from raffinose to galactose. In WT strains, Msc1 levels were higher than endogenously expressed Msc1 as early as 1 h after galactose addition (Fig S8B). These levels were even much higher after 2 h in galactose. Long-term Msc1 overexpression was toxic in both spot assays and growth curves (Fig S8C-E), and overexpressed Msc1 did not rescue WT sensitivity to phleomycin in growth curves (Fig S8D). However, in short-term experiments with Δ*msc1* cells arrested in late-M, overexpression of Msc1 for 2 h increased the proportion of cells with Rad52 foci in both HO and phleomycin (Fig 4C,D).

## Discussion

Repair of highly deleterious DSBs by HR requires an intact sister chromatid for not being mutagenic. Prior to DNA replication, when a sister is not yet available, HR is inhibited as its activity is coupled to CDK activity ^7,8^. However, in the final stages of mitosis, after sister chromatid segregation, CDK activity still remains high and thus HR remains active. In a previous work, we observed that *S. cerevisiae* resolves this paradox by approaching and coalescing the segregated sister, which could give HR one last chance to repair faithfully. We have now used a proteomic approach to search for new factors that can play a prominent role in DSB repair in late-M. From this study, we have identified the loosely characterized protein Msc1 (Fig 1A; Table S1). After validating the proteomic findings by Western blotting and microscopy (Fig 1B-E), we confirmed that the deletion mutant is more sensitive to DSBs generated in late-M (Fig 2A-D). From a mechanistic point of view, the Δ*msc1* mutant appears to affect either the formation of the presynaptic filament or homology search afterward (Fig 4E). Accordingly, HR completion is delayed compared to WT, but DSBs resection is not (Fig 3). This position downstream of the resection also fits well with the fact that the mutant has fewer Rad52 repair centers (Fig 4A-D), but at the same time has higher signs of DNA damage (hyperphosphorylated Rad53) (Fig 2E,F and S6).

Msc1 is an NE protein that belongs to the poorly-characterized Ish1 family. To date, the best characterized member of this family is Les1 from *Schizossacharomyces pombe*, which has been linked to aberrant karyokinesis ^35^. SpLes1 is a nuclear envelope protein that localizes to the bridge stalk before karyokinesis and corrals nuclear pore complexes (NPC) ^35^. Because SpLes1 and ScMsc1 are orthologs, the Msc1 deficiency could also interfere with the axis that regulates the repair of DSBs via NPCs. In *S. cerevisiae*, this axis is particularly important for DSBs that lack a nearby template ^36–39^, as is the case in late-M. In this sense, DSB targeting to the NE and the putative role of Msc1 in NPC dynamics may facilitate the search for distant templates, perhaps overcoming the barrier of the long and thin nuclear bridge ^20^.

In conclusion, through abundance proteomics we have identified a novel protein, Msc1, that enhances DSB repair in late mitosis, when the nucleus is elongated and sister chromatids have been segregated. Our data further suggest that Msc1, which is an NE protein, upregulates the assembly of Rad52 repair factories and accelerates HR completion. This work presents for the first time to the best of our knowledge a protein with a specific role in late-M HR.

## Material and Methods

### Yeast strains and experimental conditions

All yeast strains used are listed in Table S3. Strain construction was undertaken through standard transformation methods ^40^. Gene deletions and C-terminal tags were engineered using PCR methods ^41^. The MoClo Yeast Toolkit was used to create the *MSC1* overexpression plasmid following the instructions ^42^. To add *MSC1* to the system as a type 3 module, the sequence of *MSC1* with the necessary overhangs was ordered as a synthetic gene (gBlocks HiFi Gene fragments from IDT). A synonymous mutation was also added to the sequence to destroy a target for the restriction enzyme BsaI, which is used for the assembly of the different modules.

Strains were grown overnight in air orbital incubators at 25ºC in YPD media (10 g·L^−1^ yeast extract, 20 g·L^−1^ peptone and 20 g·L^−1^ glucose) unless stated otherwise. To arrest cells in late mitosis, log-phase asynchronous cultures were adjusted to OD_600_ ∼ 0.4 and the temperature was shifted to 34°C for 3 h. In most experiments, the arrested culture was split into three subcultures: one subculture was treated with phleomycin (2-10 μg·mL^−1^; Sigma-Aldrich, P9564), a second one with β-estradiol (2 μM; Sigma-Aldrich, E8875) for the induction of HO endonuclease ^43^, and the third was just treated with the vehicle (mock treatment). In general, samples were collected at the moment of the arrest and at 1 and 2 h after DNA damage. The general experimental setup is depicted in the upper branch of Fig S1. In the case experiments to analyze DNA repair (MAT switching and Rad53 inactivation), all cultures were washed twice with fresh YPD and further incubated for 4-24 h to recover from DNA damage. In resection experiments, β-estradiol was added to the culture at the late mitotic arrest and maintained for 4 h.

To synchronize cells in G2/M, 15 μg·mL^−1^ nocodazole (Nz; Sigma-Aldrich, M1404) was added and the cells held at 25°C for 3 h, with a Nz boost (7.5 μg·mL^−1^) at 2 ht. Then, G2/M cultures were treated as described above for the late mitotic arrest. When galactose induction was required, cells were grown in YP raffinose 2% (w/v) as the carbon source and galactose was added at 2% (w/v) 1 h after the DNA damage induction while keeping the yeast cultures at 34ºC.

### Proteomics

The experimental setup shown in Fig S1 was followed for comparative proteomics. After taking the corresponding samples, they were processed for mass spectrometry (MS). Firstly, samples were boiled 10 min at 80 ºC in LDS buffer with 10 mM DTT. Subsequently, proteins were separated using a pre-casted 4-12% NuPAGE Bis-Tris gel and run at 180 V for 10 min. The samples were later processed by the in-gel digestion protocol described in ^44^. In short, samples were first reduced in reduction buffer (10 mM DTT in 50 mM ammonium bicarbonate buffer (ABC buffer) at 56 ºC for 1 h and alkylated in alkylation buffer (50 mM iodoacetamide in 50 mM ABC buffer) for 45 min in the dark. Proteins were then digested overnight with 1 μg MS-grade trypsin at 37 ºC in 50 mM ABC buffer and the digested peptides were eluted onto a C18 StageTip, following the micro-purification protocol from ^45^. Each sample was measured with a 120 min method on an Exploris 480 mass spectrometer coupled to an Easy-nLC 1200 system (ThermoFisher Scientific) with a 50 cm column packed in-house with Reprosil C18. The mass spectrometer was operated with a top20 data-dependent acquisition method.

MS files were processed using the MaxQuant Software and the ENSEMBL *S*.*cerevisiae* protein database (version R64-1-1.24). The options “LFQ quantification” and “match between runs” were activated. The output files were analyzed using R. First, known contaminants, reverse hits and protein groups only identified by site were removed. Then, identified protein groups were filtered with a minimum of two quantification events per experiment. Missing values were imputed with a downshifted and compressed beta distribution within the 0.001 and 0.015 percentile of the measured values for each individual replicate. For plotting, LFQ intensities were log2 transformed. A two sample Welch t-test was performed. Volcano plots were generated by plotting -log10 (p-values) and fold changes. The threshold line for enriched proteins was defined with p-value=0.05, s=1 and c=0.5.

Finally, we used the fact that in 8 out of 24 samples we had induced the HO endonuclease to internally validate the proteomics readouts. Thus, HO was detected in 7 out of 8 HO induction experiment (HO at G2/M plus HO in late mitosis, n=4 each), whereas it was absent in all the other 16 samples (mock at G2/M, phle at G2/M, mock in late mitosis, and phle in late mitosis; n=4 each) (Supplemental Excel file 1).

### Western blots

Western blotting was carried out as reported before with minor modifications ^20^. Briefly, 5 ml samples were collected, the cell pellets were fixed in 1 mL of 20% (w/v) trichloroacetic acid TCA, and broken by vortexing for 3 min with ∼200 mg of glass beads in 2 mL tubes. Samples were then centrifuged, pellets were resuspended in 150 μL of a mixture of PAGE Laemmli Sample Buffer 1X (Bio-Rad, 1610747), Tris HCl 0.75M pH 8.0 and β-mercaptoethanol 2.5% (Sigma-Aldrich, M3148), and tubes were boiled at 95°C for 3 min and pelleted again. Total protein in the supernatant was quantified using a Qubit 4 Fluorometer (Thermo Fisher Scientific, Q33227). Proteins were resolved in general in 7.5% SDS-PAGE gels and transferred to PVFD membranes (Pall Corporation, PVM020C099). The membrane was stained with Ponceau S solution (PanReac AppliChem, A2935) as a loading reference.

The following antibodies were used for immunoblotting: The HA epitope was detected with a primary mouse monoclonal anti-HA (1:1,000; Sigma-Aldrich, H9658); the Myc epitope was detected with a primary mouse monoclonal anti-Myc (1: 5,000; Sigma-Aldrich, M4439); the Pgk1 protein was recognized with a primary mouse monoclonal anti-Pgk1 (1:5,000; Thermo Fisher Scientific, 22C5D8), the aid tag was recognized with a primary mouse monoclonal anti-miniaid (1:500; MBL, M214-3), and Rad53 was recognized with a primary mouse monoclonal anti-Rad53 (1:1000; Abcam, ab166859). A polyclonal goat anti-mouse conjugated to horseradish peroxidase (1:5,000; 1:10,000; or 1:20,000; Promega, W4021) was used as the secondary antibody. Antibodies were diluted in 5% skimmed milk TBST (TBS pH

7.5 plus 0.1% Tween 20). Proteins were detected by using the ECL reagent (GE Healthcare, RPN2232) chemiluminescence method, and visualized in a Vilber-Lourmat Fusion Solo S chamber.

Protein bands were quantified using BioProfile Bio1D software (Vilber-Lourmat) and then normalized with respect to PGK1, which was considered as the housekeeping. Subsequently, the Msc1 level detected in each type of damage was normalized with respect to its mock.

### qPCR

qPCR was performed on genomic DNA (for resection experiments) and from cDNA (for expression experiments) in 96-well 0.2ml block plates using a QuantStudio5 Real-Time PCR instrument. Reactions had a final volume of 10 μl and were prepared with PowerUp™ SYBR™ Green Master Mix (Thermo Scientific, A25741). The High Capacity RNA-to-cDNA kit (Thermo Scientific, 4387406) was used for the retrotranscription. RNA was extracted using the PureLink™ RNA Mini Kit (Thermo Scientific, 12183018A) and gDNA was extracted using glass beads/phenol Winston’s method ^46^. Each sample for resection analysis was divided into two aliquots and one aliquot was digested with the Sty-I-HF (NEB, R3500S) restriction enzyme. Primers used in the resection assay are listed in ^47^. Primers used in the expression experiments are: 5’-TGTCACCAACTGGGACGATA-3’and 5’-AACCAGCGTAAATTGGAACG-3’ for *ACT1* as control; 5’-TTGGATGACATAAAGGGTTG-3’ and 5’-GTACCTAAAATCATTCGGTG-3’ for *MSC1*.

To calculate *f*_*resected*_, it was first necessary to calculate the fraction of the extracted genomic DNA where HO had cut the MAT locus, which is simply known as *f*. For this, the following equation was used f=1-((E_HOcs_^ΔCq(t0-t)^)/(E_ADH1_^ΔCq(t0-t)^)) ^47^. Then, *f*_*resected*_, the fraction in which the resection has passed the restriction site, was calculated with this second equation f_resected_=2/(((E_RS_ ^ΔCq(digest-mock)^)/(E_ADH1_^ΔCq(digest-mock)^)+1)·f). In both cases, E is the primer efficiency and ΔC_q_ represents the difference in quantification cycles.

### Microscopy

A Zeiss Axio Observer.Z1/7 was also used. This inverted microscope was equipped with an Axiocam 702 sCMOS camera, the Colibri-7 LED excitation system, narrow-band filter cubes for covisualization of CFP, YFP/GFP, and mCherry without emission crosstalk, and a Plan-Apochromat 63x/NA 1.40 Oil M27 DIC objective.

For each field, a stack of 10-20 z-focal plane images (0.2-0.3 μm depth) was collected. In general, the images were taken from freshly harvested cells without further processing and at least 100 cells were quantified per experimental data point. The Zen Blue (Zeiss) and Fiji-ImageJ (NIH) software were used for image processing and quantification. Scale bars represent 4 μm in all cases.

For Rad52-mCherry factories three distributions were quantified: “No focus” (cells with a homogeneously diffused nuclear Rad52); “Focus” (a single Rad52 spot); and “Foci” (more than one Rad52 spot).

### Growth curves and viability analyses

For clonogenic survival assays, log-phase asynchronous cultures were adjusted to OD_600_ = 0.4 before the corresponding block and ensuing treatment. After that, 100 μL of 4:10,000 dilutions were spread onto YPD plates. Viability was measured by plotting the number of colonies grown on the plates after 3 days at 25ºC. The mock treatments yielded 400– 600 CFU/plate in these experiments.

For spot sensitivity assays cultures were grown exponentially and adjusted to an OD_600_ = 0.5 and then 5-fold serially diluted in YPD in 96-well plates. A 48-pin replica plater (Sigma-Aldrich, R2383) was used to spot ∼3 μL onto the corresponding plates, which were incubated at 25 °C for 3–4 days before taking photographs.

For growth curves, strains were first grown exponentially in YPD and then an inoculum was taken and adjusted to an initial OD_600_ = 0.05 in either fresh YPD or YPGalactose (2% w/v), without or with phleomycin (2 μg·mL^−1^). Three replicates of each culture were aliquoted in a flat-bottomed 96-well plate and real-time growth was measured in a Spark TECAN incubator by reading the OD_600_ every 15 minutes for 50 hours with shaking (96 rpm and 6mm of orbital amplitude). The mean of the three replicates was calculated to obtain the final growth curves. Two independent experiments were performed but only one is shown since the s.e.m was less than 0.1 OD_600_ for each time point.

### MAT switching assay and Southern blots

After taking the samples, genomic DNA was extracted by a lytic method. Briefly, the pellets were resuspended in 200 μl of digestion buffer (1 % SDS, 100 mM NaCl, 50 mM Tris-HCl, 10 mM EDTA and 50U Lyticase (Sigma-Aldrich, L4025)) and incubated at 37ºC. DNA was isolated by phenol:chloroform:isoamylalcohol (PanReac AppliChem, A0944), precipitated with ice-cold ethanol 100%, resuspended in TE 1X containing 10 μg.mL-1 RNase A (Roche, 10109169001) allowing the enzyme to act for a short incubation, precipitated a second time and resuspended in TE 1X. Then, the purified DNA was digested with StyI, the restriction fragments were separated on a 1.2% low EEOO LS Agarose gel, and finally Southern blotted. Southern blot was carried out by a saline downwards transference onto a positively charged nylon membrane (Hybond-N+, Amersham-GE; RPN303B) as reported before ^48^. DNA probes against *ACT1* and *MAT*a loci were synthesized using Fluorescein-12-dUTP Solution (ThermoFisher; R0101) and the Expand™ High Fidelity PCR System (Roche; 11732641001). Hybridization with fluorescein-labeled probes was performed overnight at 68°C. The next day, the membrane was incubated with an anti-fluorescein antibody coupled to alkaline phosphatase (Roche; 11426338910), and the signal was developed using CDP-star (Amersham; RPN3682) as the substrate. Detection was recorded using the VilberLourmat Fusion Solo S instrument.

For the quantification of the assays, each individual band was normalized to the *ACT1* signal corresponding to its sample. Then, a second normalization was performed for the signal of each *MAT*α band with respect to the intensity of the HO cut band after one hour of endonuclease action. Consequently, the graphs show the amount of DNA repaired by HR with respect to the total amount of DNA cut by HO.

### Data representation and statistics

Error bars in all graphs represent the standard error of the mean (s.e.m.) of independent biological replicates performed in different days. The number of replicates (n) is given in the figure legend. Graphpad Prism 9 was used for statistical tests. Differences between experimental data points were estimated using either the Mann-Whitney U test or the unpaired t-test; the test used is indicated in the figure caption. Significance is denoted by asterisks (* indicates p<0.05, ** indicates p<0.01, *** indicates p<0.001 and **** indicates p<0.0001).

In general, we used four types of graphs to represent the data: volcano plots, bar charts, marker line graphs and box plots. In box plots, the center line represents the medians, box limits represent the 25th and 75th percentile, the whiskers extend to the 5th and 95th percentiles, and the dots represent outliers.

## Data Availability

The mass spectrometry proteomics data have been deposited to the ProteomeXchange Consortium via the PRIDE partner repository with the dataset identifier PXD043515. All other data is contained within the manuscript and/or supplementary files.

## Acknowledgements

We would like to kindly thank Lorraine Symington for strains, as well as for hosting and teaching J.A-P. the MAT switching and HOcs resection methodologies. We thank the Core Facility Proteomics at the Institute of Molecular Biology (Mainz) for their support in processing and analyzing the mass-spectrometry data. We thank Mario Dejung for the development of the R script that analyzes the mass-spectrometry data.

This work was supported by the Spanish Ministry of Science and Innovation (research grants BFU2017-83954-R and PID2021-123716OB-I00 to F.M.) and the Agencia Canaria de Investigación, Innovación y Sociedad de la Información of the Canary Government (predoctoral fellowship TESIS2020010028 to S.M-S.). All grants were co-financed by the FSE Structural Funds. Part of this work was supported by the Deutsche Forschungsgemeinschaft (Project-ID 393547839-SFB 1361 to F.B.)

## Author contributions

S. M-S.: Performed all experiments shown in the main and supplementary figures and tables except for Figure 1A; constructed strains and plasmids; prepared the corresponding figures and tables (with the aid of F.M); and gave critical insights as to the direction and development of the study.

J. A-P.: Performed the experiments for the proteomics in Figure 1A; taught S.M-S. and gave critical insights as to the direction and development of the study.

L. P-M.: Performed the proteomics; analysed the proteomic data; and generated the volcano plots shown in Figure 1A.

F. B.: Performed the proteomics, analysed the proteomic data; generated the volcano plots shown in Figure 1A; was the supervisor of L.P-M.; and was responsible for funding acquisition and project administration related to proteomics.

F. M.: Supervised the project; is/was the supervisor of S.M-S. and J.A-P.; was responsible for funding acquisition and project administration related to all experiments but proteomics; gave critical insights as to the direction and development of the study; and wrote the manuscript (with the aid of S.M-S).

