## Supplemental Figures S1-S8 and tables S1-S3 for "Msc1 is a nuclear envelope protein that reinforces DNA repair in late mitosis"

### Supplemental Information

### SUPPLEMENTARY TEXT

#### On the MAT switching assay

Unlike other regions of the genome, the clean cut of the *MAT* locus by the HO endonuclease uses ectopic homologous sequences for HR repair, resulting in repair products that are discernible by Southern blot. When the sequence of the *MAT* locus determines for "a" sex (*MAT*a), the cryptic ectopic sequence *HML* is used as the preferred template so that HR leads to a gene conversion to *MAT*α (Fig 3A). Both alleles can be distinguished by a restriction length polymorphism for the enzyme *StyI*. Overall, the pattern of *StyI* recognition sequences along the alleles, together with the position of the probe, allows the identification of the uncut *MAT*a (0.9 Kb band), its HO-cut downstream product (0.7 Kb band), and the HR-driven gene conversion to *MAT*α (1.8 Kb band). The probe partially overlaps the most downstream *StyI* site of the reporter and thus recognizes other sequences downstream of the *MAT* locus. This results in another high molecular weight band (the slowest to migrate; 2.2 Kb) that is affected by neither the actual *MAT* allele nor the HO cut and can be used as an internal loading control for quantification. However, since this region is relatively close to the HO cut and could be affected by DSB resection, which could reduce the amount of signal given by the probe, we also used a second probe for a distant locus (*ACT1*, the 1.1 Kb band) as a second internal loading control.

As mentioned above, HR-driven gene conversion after the HO cut results in the appearance of the 1.8 Kb *MAT*α band. Alternatively, the DSB could also be repaired through NHEJ, restoring the *MAT*a allele (0.9 Kb band). *MAT*a could also be restored by ectopic HR if the *HMR* locus is used instead of the preferred *HML*; however, we used a strain where *HMR* has been deleted to avoid this confusing scenario. Thus, the cut *MAT*a locus can lead to only three outcomes in this *hmr*Δ strain: (i) gene conversion by HR (the HO-cut 0.7 Kb band disappears in favor of the *MAT*α 1.8 Kb band); (ii) NHEJ (the HO-cut 0.7 Kb band disappears in favor of the *MAT*a 0.9 Kb band); (iii) or the DSB remains unrepaired (neither the 1.8 Kb nor the 0.9 Kb band get enriched relative to their values at the time of the HO removal).

### SUPPLEMENTARY FIGURES AND TABLES

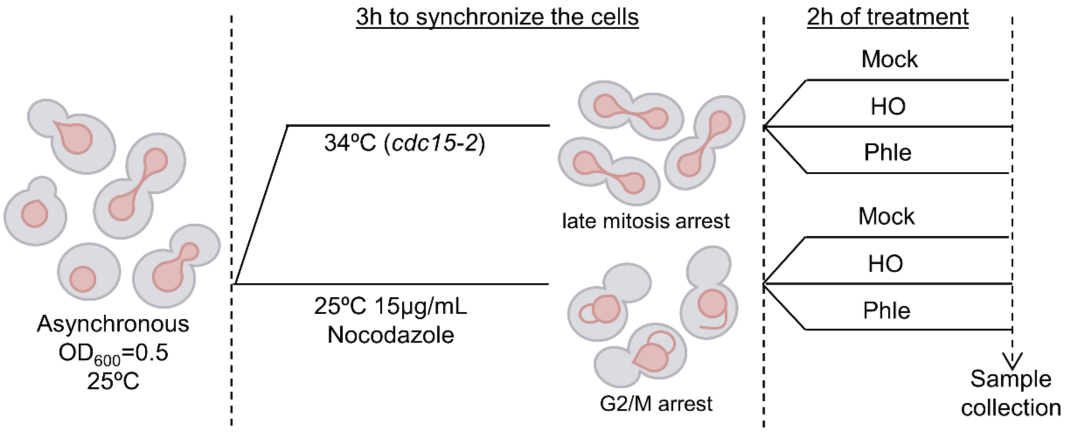


**Figure S1. Proteomics strategy for DSBs in G2/M and late mitosis. (A)** Schematic of the experimental procedure. Cells were first arrested either in G2/M by adding nocodazole or in late mitosis by incubating at 34ºC. Then, the culture was divided into three subcultures. One served as a mock control, whereas the others were treated to generate DSBs, one with β-estradiol (for HO expression) and the other with phleomycin.

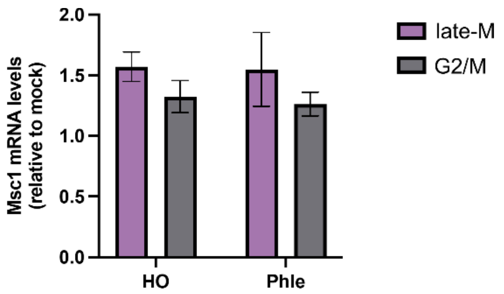


**Figure S2. Relative *MSC1* mRNA level after DSBs in G2/M and late-M.** Using the same experimental setup as for Western blots in Figure 1C, total RNA samples were collected for RT-qPCR quantification of *MSC1* mRNA levels (mean ± s.e.m., n=3).

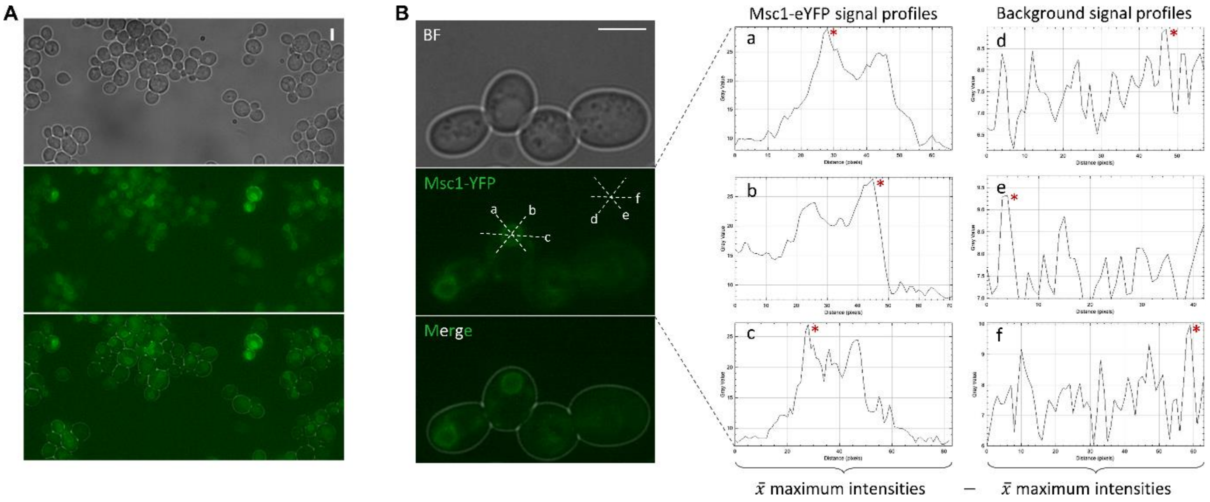


**Figure S3. Quantification of Msc1-eYFP intensity at the nuclear envelope. (A)** Representative microscopic field of asynchronous cells expressing Msc1-eYFP under its endogenous promoter. Note that Msc1 localizes to the NE and its levels are variable in the cell population. (**B**) Method for quantifying NE Msc1 levels in late-M cells. Using the ImageJ/FiJi software, each nuclear body in the elongated late-M nucleus was traversed with three selection lines converging at the center and leaving equivalent angles between the lines. Then, the maximum intensity was determined from the intensity profile when the line crossed the NE and the mean of all intensities was calculated. Background intensities were estimated in a similar way in a cell-free area. The NE and background intensities were subtracted to calculate the relative intensity of each cell. Plots of these intensities are shown in Figure 1E.

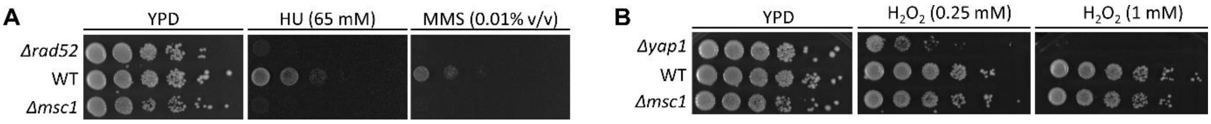


**Figure S4. Sensitivity of the *msc1*Δ mutant to replicative and oxidative stresses. (A)** Spot assays against the replicative stress agents hydroxyurea (65mM) and MMS (0.01% v/v). The first line corresponds to a *rad52*Δ mutant used as a positive control for sensitivity to DNA damage. **(B)** Spot assays against the oxidative stress agent hydrogen peroxide (0.25 and 1mM). The first line corresponds to a *yap1*Δ mutant used as a positive control for sensitivity to oxidative stress. In both cases, the different strains were grown overnight to log phase in YPD, the OD600 was adjusted to 0.5, 1:5 serial dilutions were made and then spotted onto the corresponding plates.

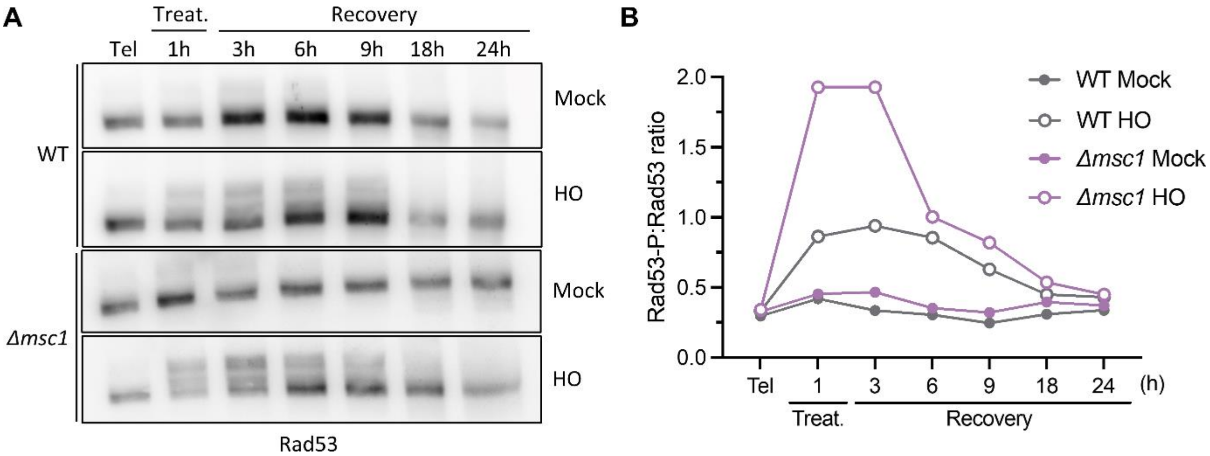


**Figure S5. Role of Msc1 in DNA damage recovery. (A)** Western blot of the loss of Rad53 hyperphosphorylation after DSB recovery in the WT and the *msc1*Δ strains. The experimental setup was as is Figure S1, but only late-M HO-mediated DSBs were generated with 2 μM BE. After 1h, BE was washed off and samples were collected after 3, 6, 9, 18 and 24 h. **(B)** Quantification of the Rad53-P:Rad53 ratio through the time course.

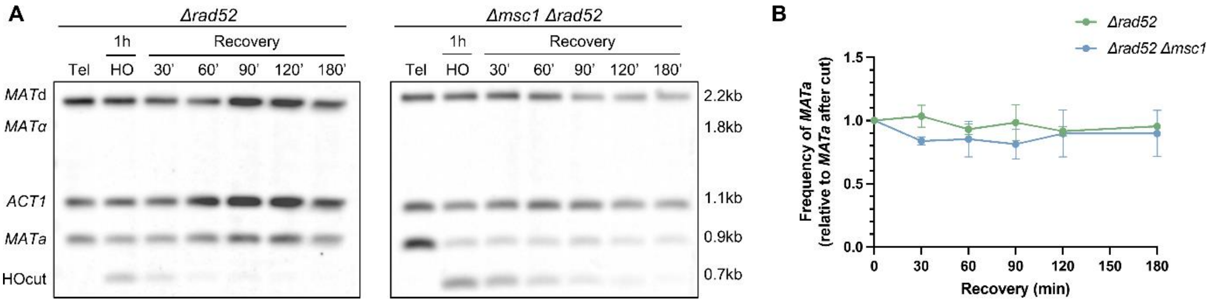


**Figure S6. Late mitotic repair of the HO-mediated DSB entirely depends on HR.** Related to Figure 3A-C. **(A)** Representative Southern blots of the MAT switching assay in the *MSC1 rad52*Δ and *msc1*Δ *rad52*Δ strains. Note that the *MAT*α HR product is not obtained. (**B**) Quantification of the *MAT*a band through the experiment (mean ± s.e.m., n=3). The band was normalized to the amount of the *MAT*a that remained after HO induction. Note that even in these *rad52*Δ strains, unable to repair by HR (i.e., no *MAT*α band), no signs of NHEJ are seen either (*MAT*a increase during the DSB recovery).

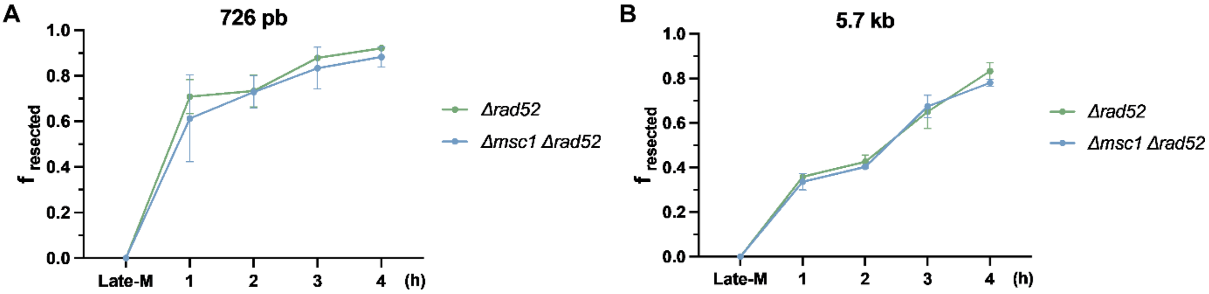


**Figure S7. Late mitotic resection of the HO-mediated DSB ends in the Δ*rad52* and the Δ*rad52* Δ*msc1* strains.** Related to Fig 3E. On the left, resection kinetics for an amplicon located 726 bp downstream of the HO-generated DSB (mean ± s.e.m., n=2). On the right, resection kinetics for an amplicon located 5.7 bp downstream of the HO-generated DSB (mean ± s.e.m., n=2); f resected is the fraction of resected DNA.

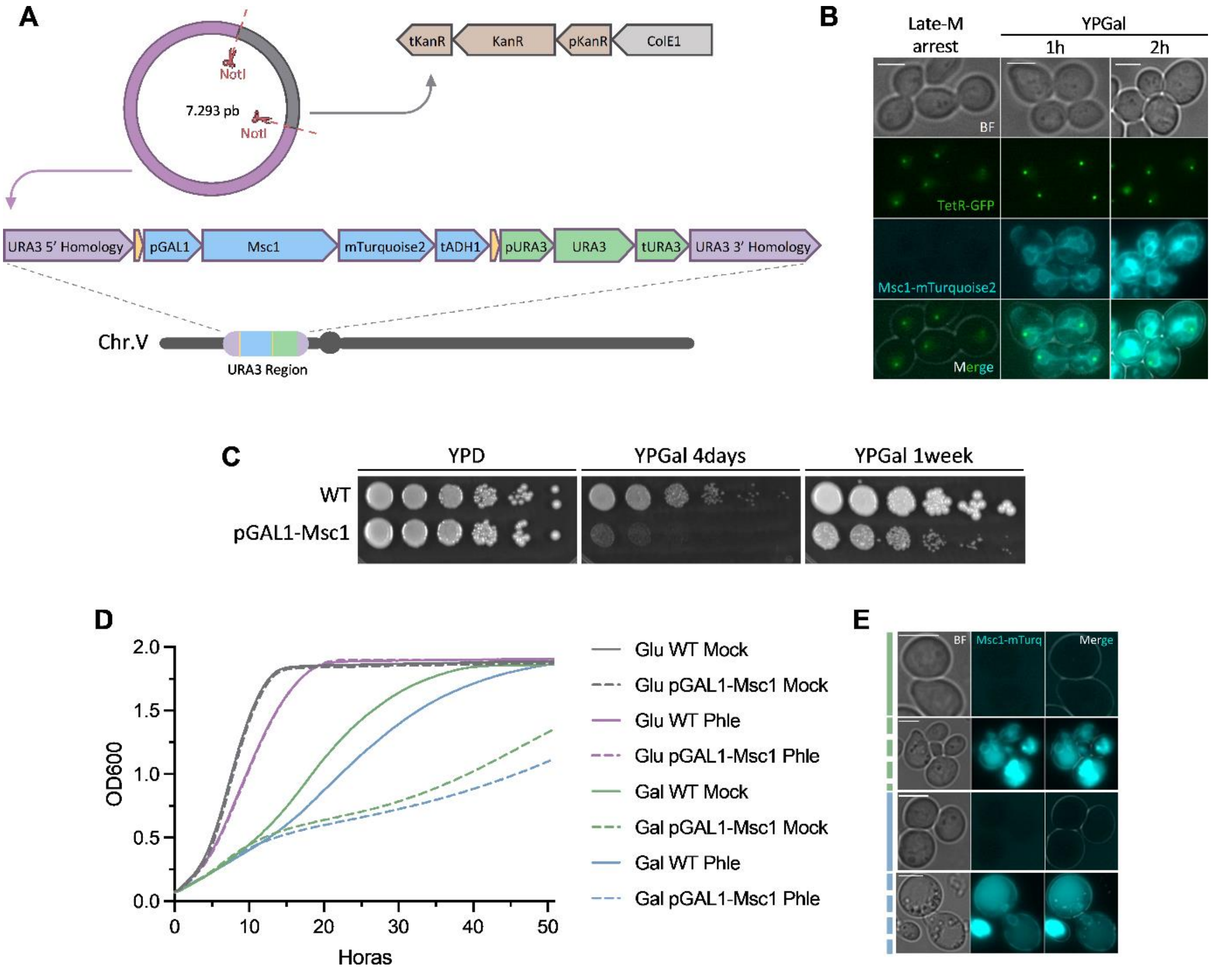


**Figure S8. Short- and long-term Msc1 overexpression profiles. (A)** Schematics of the integrative plasmid created by modular cloning for overexpressing *MSC1* under the *GAL* promoter. The construct is designed to be integrated ectopically at the *URA3* locus. Msc1 includes the mTurquoise2 tag at the C-terminus to verify expression by microscopy. Another variant of this plasmid was also created with the only difference that their integration is at the *HO* locus and has *LEU2* as marker. **(B)** Confirmation that Msc1-mTurquoise2 is overexpressed in a late-M arrest after a short incubation in YP galactose. Msc1-mTurquoise2 was clearly detected with less exposure time and intensity from the fluorescence source than the endogenous Msc1-eYFP after just 1h incubation; after 2h, signal saturation was common. Overexpressed Msc1 was located at other peripheral structures (ER and/or plasma membrane) aside from the NE. **(C)** Spot assay of long-term Msc1 overexpression. Parental (WT) and the *pGAL-MSC1* strains were grown overnight in YPD to log phase, the $OD_{600}$ was adjusted to 0.5, 1:5 serial dilutions were made and then spotted onto the corresponding plates. The YPGal plate was photographed after 4 and 7 days. Note that Msc1 overexpression is toxic. **(D)** Growth curves of the WT and *pGAL-MSC1* strains under different inducing conditions (Glu or Gal) and with (Phle) or without (mock) concomitant moderate DNA damage caused by phleomycin (2 $\mu g \cdot mL^{-1}$). The initial inoculum for both strains was set to $OD_{600}$=0.05. Data are the mean of two independent experiments (the s.e.m is not represented but was less than 0.1 throughout). Note that both strains grow to the same extent with glucose as the carbon source, regardless of whether phleomycin is added or not. With galactose as the carbon source, the growth of the strain with galactose-dependent Msc1 overexpression is clearly slowed. Msc1 overexpression neither increases nor suppresses Phle sensitivity. **(E)**

Microscopy of cells taken from the growth curves in YPGal after 50h. In WT there is no signal in the blue channel, while in strains with the *pGAL1-MSC1-mTurquoise2* system a strong yet mislocated Msc1 signal was observed.

**Table S1. Genes that encode proteins that significantly change their levels upon DSBs in late-M but not in G2/M.**

| HO-mediated DSBs | | |
|---|---|---|
| Up-regulated | | Down-regulated |
| *YNL333W* (*SNZ2*) | *YDR221W (GTB1)* | *YHR083W (SAM35)* |
| *YFL059W (SNZ3)* | *YPL214C (THI6)* | *YLR417W (VPS36)* |
| ***YLR178C (TFS1)**** | *YGR281W (YOR1)* | *YDR440W (DOT1)* |
| ***YPR160W (GPH1)**** | ***YML128C (MSC1)**** | ***YEL046C (GLY1)**** |
| ***YGR256W (GND2)**** | *YPR172W* | *YKL082C (RRP14)* |
| *YOL036W* | *YMR271C (URA10)* | *YKR020W (VPS51)* |
| ***YPR184W (GDB1)**** | | |
| Phleomycin-mediated DSBs | | |
| Up-regulated | | Down-regulated |
| *YKL107W* | *YNR034W-A (EGO4)* | *YBR014C (GRX7)* |
| *YMR105C (PGM2)* | *YOR173W (DCS2)* | ***YEL046C (GLY1)**** |
| ***YGR256W (GND2)**** | *YDL204W (RTN2)* | *YDR246W (TRS23)* |
| *YMR196W* | *YFL014W (HSP12)* | *YGR283C (UPA1)* |
| ***YLR178C (TFS1)**** | *YEL039C (CYC7)* | *YER126C (NSA2)* |
| ***YML128C (MSC1)**** | *YDR345C (HXT3)* | *YLR056W (ERG3)* |
| *YLR001C* | *YFL011W (HXT10)* | |
| ***YPR184W (GDB1)**** | *YOL156W (HXT11)* | |
| *YIL136W (OM45)* | *YJL219W (HXT9)* | |
| *YLR258W (GSY2)* | *YGR248W (SOL4)* | |
| *YGR043C (NQM1)* | ***YPR160W (GPH1)**** | |
| *YBR072W (HSP26)* | *YOR120W (GCY1)* | |
| *YBR234C (ARC40)* | | |

*Genes whose products change significantly against both HO-mediated and phleomycin-mediated DNA damage are highlighted in bold type.

**Table S2. Changes of abundance (log2) of known DSB signaling and repair proteins upon DSBs in G2/M and late-M.**

| Protein[1] | G2/M with HO | | G2/M with Phle | | Late-M with HO | | Late-M with Phle | |
|---|---|---|---|---|---|---|---|---|
| | $-\log_{10}$ (p-value) | $\log_2$ (fold change) | $-\log_{10}$ (p-value) | $\log_2$ (fold change) | $-\log_{10}$ (p-value) | $\log_2$ (fold change) | $-\log_{10}$ (p-value) | $\log_2$ (fold change) |
| Chk1 | 0,47 | -0,17 | 0,10 | -0,34 | 1,33 | -0,35 | 0,53 | -0,12 |
| Dcc1 | - | - | - | - | 0,18 | 0,02 | 1,08 | 0,51 |
| Ddc1 | - | - | - | - | - | - | - | - |
| Ddc2 | 0,56 | -0,14 | - | - | 0,15 | -0,08 | 0,20 | -0,12 |
| Dna2 | - | - | 2,03 | 0,25 | - | - | - | - |
| Dnl4 | - | - | - | - | - | - | - | - |
| Exo1 | - | - | - | - | - | - | - | - |
| Mec1 | - | - | 0,61 | -0,77 | - | - | 0,07 | -0,43 |
| Mms4 | - | - | - | - | - | - | - | - |
| Mph1 | 0,29 | 0,01 | - | - | - | - | - | - |
| Mre11 | 0.21 | 0.37 | - | - | 0.20 | 0.14 | 0.50 | 0.39 |
| Msh2 | 0,10 | 0,08 | 0,27 | -0,11 | 0,26 | -0,21 | 0,06 | -0,47 |
| Mus81 | - | - | - | - | - | - | - | - |
| Rad5 | - | - | - | - | - | - | - | - |
| Rad9 | - | - | - | - | - | - | - | - |
| Rad50 | 0.94 | 0.52 | 0.71 | 0.73 | 0.76 | 0.38 | 1.98 | 0.47 |
| Rad51 | 0.86 | 0.31 | 1.34 | 0.62 | 0.02 | -0.01 | 0.68 | 0.51 |
| Rad52 | 0.44 | 0.15 | 0.27 | 0.15 | 0.11 | -0.54 | 0.33 | -0.19 |
| Rad53 | 0.33 | -0.01 | - | - | 0.15 | 0.24 | 0.06 | 0.13 |
| Rad54 | - | - | - | - | - | - | - | - |
| Rad55 | - | - | - | - | - | - | - | - |
| Rad57 | - | - | - | - | - | - | - | - |
| Rad59 | - | - | - | - | - | - | 1,09 | -0,94 |
| Rfa1 | 0,08 | 0,01 | 0,01 | 0,01 | 1,80 | -0,39 | 2,91 | -0,48 |
| Rmi1 | - | - | - | - | - | - | - | - |
| Sae2 | - | - | - | - | - | - | - | - |
| Sgs1 | - | - | - | - | - | - | - | - |
| Slx1 | - | - | - | - | - | - | - | - |
| Slx4 | - | - | - | - | - | - | - | - |
| Smc1 | 1,58 | 0,38 | 2,10 | 0,41 | 0,02 | -0,01 | 0,46 | 0,27 |
| Smc2 | 0,63 | 1,09 | 0,98 | 0,16 | 0,09 | 0,40 | 0,69 | 0,62 |
| Smc3 | 1,95 | 0,63 | 1,07 | 0,32 | 0,19 | -0,09 | 0,04 | 0,02 |
| Smc4 | 0,52 | 0,51 | 0,14 | 0,31 | 0,76 | 0,47 | 0,62 | 0,22 |
| Smc5 | - | - | - | - | - | - | 1,48 | 0,18 |
| Smc6 | - | - | - | - | - | - | - | - |
| Srs2 | - | - | - | - | - | - | - | - |
| Tel1 | - | - | - | - | - | - | - | - |
| Top3 | - | - | - | - | - | - | - | - |
| Xrs2 | - | - | - | - | - | - | - | - |

| | | | | | | | | |
|---|---|---|---|---|---|---|---|---|
| Yen1 | - | - | - | - | - | - | - | - |
| Yku70 | - | - | - | - | - | - | - | - |
| Yku80 | - | - | - | - | 0.19 | -0.06 | 0.29 | 0.17 |

[1] Proteins are ordered alphabetically. “-” indicates that the protein was not detected by the inclusion criteria set for the proteomics analysis (see material and methods).

**Table S3. Strains used in this work.**

| Name | Genotype[1] | Origin | Use |
|---|---|---|---|
| FM2531 | *MATa bar1Δ leu2-3,112 ura3-52 his3-Δ200 trp1-Δ63 ade2-1 lys2-801; cdc15-2:9myc::Hph; HMLα Δhmr::HIS3MX; leu2-3::LexA-TF-PlexOp:HO::LEU2* | Machín lab | Figs 1A; 2; 3; S1; S2; S4; S5<br>Table S1; S2 |
| FM2790 | FM2531; *MSC1:6HA::KanMX* | This Study | Figs 1B,C |
| FM2831 | FM2531; *MSC1:eYFP::KanMX* | This Study | Figs 1D,E; S3 |
| FM82 | *MATa bar1Δ leu2-3,112 ura3-52 his3-Δ200 trp1-Δ63 ade2-1 lys2-801; Δrad52::kanMX4* | Machín lab | Figs 2A,B |
| FM2808 | FM2531; *Δmsc1::NatNT2* | This Study | Figs 2; 3; S4 |
| FM2929 | FM2531; *RAD52:mCherry::NatNT2* | This Study | Figs 4A,B |
| FM2891 | FM2808; *RAD52:mCherry::KanMX* | This Study | Figs 4A,B |
| FM3071 | FM2891; *NUP49:eGFP:TRP1; ura3-52::GAL1p:MSC1:mTurquoise2::URA3* | This study | Fig 4C,D |
| FM2915 | FM2808; *Δrad52::KanMX* | This Study | Figs S6; S7 |
| FM2917 | FM2531; *Δrad52::KanMX* | This Study | Figs S6; S7 |
| FM2317 | *MATa bar1Δ leu2-3,112 ura3-52 his3-Δ200 trp1-Δ63 ade2-1 lys2-801; ade2-1::TetR:YFP::ADE2; tetO(5.6Kb)::194Kb-ChrXII::HIS3; cdc15-2:9myc::Hph; CIN8:mCherry::KanMX6* | Machín lab | Fig S8C-E |
| FM2817 | FM2317; *ho::GAL1p:MSC1:Turquoise::LEU2::ho* | This Study | Fig S8 |
| FM630 | *MATa his3Δ1 leu2Δ0 met15Δ0 ura3Δ0 Δyap1::kanMX4* | Euroscarf | Fig S4B |

[1] Semicolon (“;”) separates genetic modifications accomplished sequentially through transformation. Intermediate strains are omitted.
